# A/T/N polygenic risk score for cognitive decline in old age

**DOI:** 10.1101/838847

**Authors:** Annah M. Moore, Teresa J. Filshtein, Logan Dumitrescu, Amal Harrati, Fanny Elahi, Elizabeth C. Mormino, Yuetiva Deming, Brian W. Kunkle, Dan M. Mungas, Trey Hedden, Liana G. Apostolova, Andrew J. Saykin, Danai Chasioti, Qiongshi Lu, Jessica Dennis, Julia Sealock, Lea K. Davis, David W. Fardo, Rachel Buckley, Timothy J. Hohman

## Abstract

**INTRODUCTION:** We developed a novel polygenic risk score (PRS) based on the A/T/N (amyloid plaques (A), phosphorylated tau tangles (T), and neurodegeneration (N)) framework and compared a PRS based on clinical AD diagnosis to assess which was a better predictor of cognitive decline.

**METHODS:** We used summary statistics from genome wide association studies of cerebrospinal fluid amyloid-β (Aβ_42_) and phosphorylated-tau (ptau_181_), left hippocampal volume (LHIPV), and late-onset AD dementia to calculate PRS for 1181 participants in the Alzheimer’s Disease Neuroimaging Initiative (ADNI). Individual PRS were averaged to generate a composite A/T/N PRS. We assessed the association of PRS with baseline and longitudinal cognitive composites of executive function and memory.

**RESULTS:** The A/T/N PRS showed superior predictive performance on AD biomarkers and executive function decline compared to the clinical AD PRS.

**DISCUSSION:** Results suggest that integration of genetic risk across AD biomarkers may improve prediction of disease progression.

**Research in Context:** *Systematic Review:* Authors reviewed relevant literature using PubMed and Google Scholar. Key studies that generated and validated polygenic risk scores (PRS) for clinical and pathologic AD were cited. PRS scores have been increasingly used in the literature but clinical utility continues to be questioned.

*Interpretation:* In the current research landscape concerning PRS clinical utility in the AD space, there is room for model improvement and our hypothesis was that a PRS with integrated risk for AD biomarkers could yield a better model for cognitive decline.

*Future Directions:* This study serves as proof-of-concept that encourages future study of integrated PRS across disease markers and utility in taking an A/T/N (amyloidosis, tauopathy and neurodegeneration) focused approach to genetic risk for cognitive decline and AD.

## 1. Introduction

Alzheimer’s disease (AD) currently affects roughly 5.7 million people in the United States[1]. By 2025, it is expected that 7.1 million people will be affected [1], highlighting the urgency for progress in AD-focused research. AD is highly heritable, with as much as 79% shared heritable risk reported in twin studies [2], although the genetic architecture is complex including notable polygenicity. Phenotypically, AD includes a long prodrome in which neuropathology begins to accumulate decades before cognitive symptoms can be detected [3, 4]. For that reason, early detection is critical to prevent progression of the disease [5, 6]. A combined screening approach that integrates biomarker, genomic, and clinical information will likely be required to optimize early identification and prevention of AD progression [7, 8].

Polygenic risk scores (PRS) represent a tool for early AD risk detection; however, studies that used PRS to predict AD case/control status reported predictive accuracy (AUC_max_) of less than 83% [9-12], and the clinical utility of PRS beyond the *APOE* locus have been questioned [12, 13]. A wealth of genetic data from genome-wide association studies (GWAS) have become available in recent years [14], and incorporation of additional genetic data may represent a strategy to improve PRS predictive ability. In 2010, the largest GWAS of AD involved ∼16,000 participants [15], compared to the recently completed AD GWAS that included ∼600,000 participants [16]. This increase in GWAS data have enabled an increase in the number of single nucleotide polymorphisms (SNPs) used to calculate AD genetic risk scores. Some of the first PRS studies used as few as 5 SNPs to calculate AD risk, compared to 205,068 SNPs in a more recent study [17]. Despite the progress made in genomic studies of AD, PRS continue to show limited power, and innovative analytical strategies from novel perspectives are required if PRS are going to attain clinical utility.

Improvement for PRS may come from GWAS of AD endophenotypes which focus on neuropathological features of the disease. In fact, *in vivo* biomarkers of AD neuropathology have become a central feature used to identify cognitively normal individuals who are at the greatest risk of cognitive decline [4, 18]. Additionally, the National Institute on Aging and Alzheimer’s Association released a research framework for AD in which the pathological accumulation of amyloid plaques (A), neurofibrillary tangles composed of tau (T), and neurodegeneration (N) are recommended to be included in diagnostic categories of AD used for research [19]. While this A/T/N framework has emerged in studies of preclinical disease, it has not been integrated into PRS for AD despite the availability of GWAS summary statistics for autopsy and *in vivo* measures of A/T/N [20, 21]. Indeed, a previous study suggested that shared genetic drivers for hippocampal volume (a marker of neurodegeneration) and AD may exist [22]. Thus, integrating the common and independent genetic drivers of clinical AD and A/T/N could produce models with higher predictive capacity for cognitive decline in late life.

We set out to determine how PRS for CSF biomarkers of AD (amyloid and ptau_181_), hippocampal volume, and clinical AD relate to biomarkers of AD and longitudinal cognitive performance. We then developed a novel composite PRS using the A/T/N framework to integrate genetic risk for AD biomarkers, hippocampal volume, and clinical AD diagnosis into a single score (A/T/N PRS). We compared the predictive capabilities of the A/T/N score to a PRS score for clinical AD and hypothesized that the A/T/N PRS would serve as a more predictive genetic risk profile compared to a PRS for clinical AD alone.

## 2. Methods

### 2.1 Participants

Participants were drawn from the Alzheimer’s Disease Neuroimaging Initiative database (ADNI; adni.loni.usc.edu) launched in 2003 as a public-private partnership. The ADNI, ADNI-GO and ADNI-2 studies enrolled more than 1,500 participants, aged 55–90 years, excluding serious neurological disease, other than AD, and history of brain lesion, head trauma, or psychoactive medication use (for full inclusion/exclusion criteria see http://www.adni-info.org). Written informed consent was obtained from all participants at each site, and analysis of ADNI’s publicly available database was approved by our local Institutional Review Board prior to data analysis.

We accessed publicly available participant data from ADNI on July 12, 2018. ADNI enrollment criteria are outlined in the ADNI protocol (http://www.adniinfo.org/Scientists/AboutADNI.aspx). For the cognitive analyses, we included all participants who had genomic data and longitudinal cognitive (memory and executive function) data, yielding a sample size of 1,181 participants. From this sample, 1,086 subjects also had brain MRI data and were included in neuroimaging analyses. CSF biomarker data was available for 826 participants of those whom had genomic and cognitive data. Demographics of participants with genetic and cognitive data are shown in **Table 1**.

**Table 1.** Cohort demographics and summary statistics

| Participants with genetic and cognitive data | Clinical Diagnosis |  |  | Total (N=1182) | P |
| --- | --- | --- | --- | --- | --- |
|  | Normal Cognition (n=336) | Mild Cognitive Impairment (n=640) | Alzheimer's Disease (n=205) |  |  |
| Age at baseline, years | 75.3 ± 5.4 | 73.6 ± 7.5 | 75.6 ± 7.8 | 74.4 ± 7.1 | 0.72 |
| Female, no. (%) | 161 (48) | 250 (39) | 88 (43) | 499 (42) | **0.03** |
| Education, years | 16.3 ± 2.7 | 15.9 ± 2.9 | 15.0 ± 3.0 | 15.9 ± 2.9 | **7.4E-7** |
| *APOE*-ε4 carrier | 90 (27%) | 313 (49%) | 138 (67%) | 541 (46%) | **6.1E-24** |
| Baseline Memory | 1.0 ± 0.5 | 0.1 ± 0.7 | -0.9 ± 0.6 | 0.2 ± 0.9 | **7.9E-178** |
| Baseline Executive Function | 0.8 ± 0.5 | 0.2 ± 0.8 | -0.9 ± 0.6 | 0.2 ± 1.0 | **1.2E-97** |

Values are presented as mean±standard deviation, unless otherwise indicated. **Boldface** indicates $P < 0.05$ .

### 2.2 CSF collection and assays for Aβ_42_ and ptau_181_

The ADNI protocol for CSF collection and quantification of Aβ_42_ and ptau_181_ biomarkers has been detailed previously, and used the multiplex xMAP Luminex platform [23, 24]. For this study, the UPenn master data set that was available on the ADNI website was downloaded. The first biomarker measurement for each participant was used as a continuous variable in statistical models.

### 2.3 Diagnostic criteria

Full details of these diagnostic criteria have been previously published [25]. Briefly, Normal Cognition (NC) participants had a Mini-Mental State Exam [26] (MMSE) score between 24-30 (inclusive), a Clinical Dementia Rating (CDR) of 0, and were non-depressed. Mild Cognitive Impairment (MCI) participants scored between 24-30 (inclusive) on the MMSE, had a memory complaint or objective memory loss as measured by the Wechsler Memory Scale-Revised (WMS-R) Logical Memory II, a CDR of 0.5, and absence of impairments significant enough to fit criteria for dementia. An AD diagnosis required MMSE scores between 20-26 (inclusive), a CDR score of 0.5-1.0 and meeting probable AD criteria [27].

### 2.4 Brain Imaging

The ADNI neuroimaging protocol has been reported in detail elsewhere [28]. Images for the current study included original uncorrected 1.5T (ADNI-1) and 3.0T (ADNI-2, ADNI-GO) T1-weighted high-resolution three-dimensional structural data. Cortical reconstruction and volumetric segmentation were performed with the FreeSurfer image analysis suite version 4.3 in ADNI-1 and 5.1 in ADNI-2 [29-31] (http://surfer.nmr.mgh.harvard.edu/). FreeSurfer processing in ADNI has been described in detail elsewhere [32]. FreeSurfer scans were assigned a quality control (QC) value of pass, fail or partial [33]. We excluded all scans that did not have a pass QC value. We used left hippocampal volume as our primary neuroimaging outcome measurement and included a measurement of intracranial volume (ICV) and scanner strength as covariates in all volumetric analyses, with volumetric measurements defined in FreeSurfer [34].

### 2.5 Neuropsychological Testing

The ADNI neuropsychological protocol, including calculation of episodic memory and executive function composite measures, has been reported previously [35, 36]. Memory (ADNI-MEM) and executive function (ADNI-EF) composite scores were used for this study. ADNI-MEM included a composite z-score based on item-level data from Rey Auditory Verbal Learning Test, Mini-Mental State Examination (MMSE), AD Assessment Scale-Cognitive Test, and Logical Memory I and II. ADNI-EF included item-level data from Trail Making Test Parts A and B, Digit Symbol, Digit Span Backward, Animal Fluency, Vegetable Fluency, and Clock Drawing Test.

#### Genotyping and Genetic Quality Control Procedures

ADNI genotyping was performed using the Illumina Human610-Quad BeadChip (ADNI-1), the HumanOmniExpress BeadChip (ADNI-GO/2), or the Illumina Omni 2.5M WGS platform (Illumina, Inc., San Diego, CA). Quality control was performed using PLINK [37] (http://pngu.mgh.harvard.edu/purcell/plink) with a 95% threshold for genotyping efficiency applied and a minimum minor allele frequency of 0.01. SNPs outside of Hardy-Weinberg equilibrium (p<1×10^6^) were removed. Participants were excluded if they had a call rate <99%, if there was an inconsistency between reported and genetic sex, or if relatedness to another sample was established (Pihat > 0.4), in which case one participant was selected to remain in the dataset at random (9 samples removed). Imputation was performed on the Michigan Imputation Server (https://imputationserver.sph.umich.edu/index.html) using the HRC r1.1.2016 reference panel. Population structure was analyzed using the fastStructure software package [38].

### 2.6 Polygenic Risk Score Calculation

We utilized data from a GWAS for CSF biomarkers published by Deming and colleagues [20]. The original data for this study was collected from 3,146 individuals across nine cohort studies, including ADNI. After removing ADNI participants [N=390 (ADNI-1), 397 (ADNI-2)], the GWAS was re-run using data from 2,359 participants in the remaining seven cohorts. Summary results of the re-analyses excluding ADNI participants are available in Supplementary Materials.

A GWAS for clinical AD using data from the International Genomics of Alzheimer’s Project (IGAP)[39, 40] was re-run excluding 441 ADNI participants (55,931 participants remained across the Alzheimer’s Disease Genetic Consortium (ADGC), the Cohorts for Heart and Aging Research in Genomic Epidemiology (CHARGE) Consortium, the European Alzheimer’s Disease Initiative (EADI) and the Genetic and Environmental Risk in Alzheimer’s Disease (GERAD) Consortium). Summary results of the re-analysis are presented in Supplementary Materials.

Publicly available summary statistics from a UK Biobank GWAS of brain imaging phenotypes that included 8,428 participants [21] was utilized to generate ADNI participant PRS for left hippocampal volume.

PRS were calculated for each outcome in PLINK 1.9 using the scoring function (https://www.cog-genomics.org/plink/1.9/) based on summary statistics from overlapping SNPs in the ADNI cohort and the GWAS studies detailed above. The number of overlapping SNPs to calculate PRS are shown in **Supplemental Table 1**. We calculated PRS scores using a linkage disequilibrium threshold of 0.25 and physical distance of 200kb for clumping to select independent SNPs. A significance threshold of 0.01 was used for index SNPs, with a secondary significance threshold of 0.5 for clumped SNPs. Scores were generated including and excluding the *APOE* region, defined as 1MB up and downstream of the gene (chromosome 19, position 44,409,039 to 46,412,650). The PRS for CSF Aβ_42_ was multiplied by −1 to align a higher risk score with decreased CSF Aβ_42_ (and increased brain Aβ_42_ concentration). Scores were rank inverse-normal transformed to place them on the same scale [41]. Individual scores were averaged to generate a composite A/T/N score.

All studies were approved by the institutional review board (IRB) of each participating location. Data sharing was carried out within the guidelines of IRB protocols.

### 2.7 Statistical Analyses

Data were analyzed using R (version 3.5.1, https://www.r-project.org). All associations were run on each of the four individual PRS (ie, clinical AD, Aβ_42_, ptau, and LHIPV) and on the combined A/T/N PRS. Linear regression models assessing PRS score associations with CSF biomarkers (ptau_181_, Aβ_42_) covaried for age, sex, years of education, and CSF measurement batch. Linear regression models that evaluated PRS associations with left hippocampal volume (LHIPV) covaried for age, sex, years of education, baseline intracranial volume, and strength of magnet used for brain image acquisition. A binary logistic regression model assessed PRS associations with last available diagnosis (AD compared to NC, MCI diagnosed participants were excluded), and covaried for age, sex, and years of education.

We also assessed associations between PRS and baseline and longitudinal cognitive performance. A linear regression model covaried for age, sex, and years of education assessed associations between PRS and baseline cognition (memory and executive function). A mixed effects linear regression model assessed associations between PRS and longitudinal cognition (memory and executive function). Fixed effects in the model included age, sex, years of education and an interaction term for PRS × interval representing years between visits and baseline. Random effects included the intercept and interval term (slope expressed as years from the baseline visit). The pseudo R^2^ reported was the marginal coefficient of determination associated with the fixed effects of the model. A Bonferroni correction was applied to account for all 40 models tested in this study (8 outcomes × 5 PRS scores) and results that remained statistically significant after correction are indicated.

PRS performance was measured using p-value significance and adjusted R^2^ values to compare the variance explained across PRS and overall models. For mixed effects models, the marginal coefficient of determination is reported as pseudo R^2^. Sensitivity analyses were performed excluding individuals diagnosed with clinical AD or excluding clinically diagnosed AD and MCI participants. Additionally, all PRS were generated including and excluding the *APOE* region. Sex interactions on all outcomes were also investigated, and if overall interactions were found, sex-stratified analyses were performed.

## 3. Results

### 3.1 PRS Validation

The predictive performance of PRS on CSF Aβ_42_ were compared. Not surprisingly, the CSF Aβ_42_ PRS was negatively associated with measured CSF Aβ_42_ levels, indicating a higher CSF Aβ_42_ PRS is associated with higher brain Aβ_42_ burden. The CSF Aβ_42_ PRS model explained 4.6% of the variance in CSF Aβ_42_ levels (**Supplemental Table 2**, p=4.1E-06). The association remained significant after removing the *APOE* region (p=6.5E-04, R^2^=0.035) and when restricting the sample to cognitively normal and MCI individuals (p=3.7E-05, R^2^=0.045). Notably, the A/T/N score was the best predictor of CSF Aβ_42_, with the model explaining 7.6% of the variance (**Supplemental Figure 1**, p=6.8E-12) and this score was predictive of CSF Aβ_42_ in normal cognition controls (p=1.1E-03, R^2^=0.076).

CSF Ptau PRS showed a significant positive association with CSF ptau_181_ concentrations (**Supplemental Table 3**, R^2^=0.038, p=0.003), however this association did not meet the Bonferroni threshold for all 40 models tested. This association remained significant after excluding the *APOE* region (p=0.01, R^2^=0.046) and after exclusion of AD participants (p=1.1E-03, R^2^=0.059). Again, the A/T/N PRS showed the strongest association with CSF ptau_181_ concentrations compared to other scores, where a higher A/T/N PRS was associated with higher CSF ptau levels (p=8.3E-06, R^2^=0.051).

We also validated the hippocampal volume PRS, whereby a higher PRS for left hippocampal volume was associated with greater left hippocampal volume at baseline and the model explained 22.2% of variance in the LHIPV data (**Supplemental Table 4**, p=8.9E-04). Notably, this association was significant after excluding the *APOE* region. The PRS for clinical AD (calculated using a subset of the original IGAP data), and CSF Aβ_42_ PRS were also significantly associated with baseline left hippocampal volume. The clinical AD PRS score showed the strongest association and accounted for the highest variance in this outcome (R^2^=0.223, p=3.0E-04), although all PRS had similar effect sizes (R^2^ ranging from 0.215 to 0.223).

As expected, the clinical AD PRS showed the strongest association with AD diagnosis (**Supplemental Table 5**, p=7.4E-08, R^2^=0.111) and remained significant after exclusion of the *APOE* region (p=1.3E-03, R^2^=0.079). The ATN and CSF Aβ_42_ PRS were also significantly associated with AD diagnosis and remained significant after *APOE* region exclusion, however the CSF Aβ_42_ PRS association did not remain significant after Bonferroni correction.

### 3.2 PRS Association with Baseline Cognition

Higher A/T/N PRS and clinical AD PRS were significantly associated with lower baseline executive function in ADNI (**Table 2, Supplemental Figure 2**; A/T/N p=0.004, R^2^=0.115; Clinical AD p=0.02, R^2^=0.112), and these PRS models accounted for 11.2-11.4% of variability in baseline executive function but did not meet the Bonferroni correction threshold. These associations were not significant after exclusion of the *APOE* region but remained significant after exclusion of AD participants. It is noteworthy that the CSF Aβ_42_ PRS model also showed a nominal association with baseline executive function (p=0.04).

**Table 2.** PRS associations with baseline executive function performance.

|  |  |  |  | No *APOE* | NC/MCI | NC |
| --- | --- | --- | --- | --- | --- | --- |
|  |  |  |  |  | N = 845 | N = 336 |
| PRS Score | Beta | P | Adj R<sup>2</sup> | P | P | P |
| **A/T/N** | -0.14 | **0.004** | 0.115 | 0.100 | **1.56E-04\*** | 0.187 |
| **Clinical AD** | -0.06 | **0.019** | 0.112 | 0.602 | **7.81E-04\*** | 0.472 |
| **CSF AB<sub>42</sub>** | -0.05 | **0.044** | 0.111 | 0.187 | 0.066 | 0.524 |
| CSF Ptau | -0.05 | 0.088 | 0.110 | 0.168 | 0.064 | 0.220 |
| LHIPV | -0.01 | 0.763 | 0.108 | 0.769 | 0.210 | 0.767 |

**Boldface** indicates $P < 0.05$ . \* $P < 1.25E-03$ Bonferroni threshold. N=1,182 unless noted otherwise. No *APOE* denotes PRS excluding *APOE* region results. NC/MCI and NC indicate sensitivity analysis results with the denoted diagnostic groups (as assessed at baseline).

The clinical AD and A/T/N PRS were associated with baseline memory performance (**Table 3, Supplemental Figure 3**; A/T/N p=0.048, R^2^=0.093; Clinical AD p=6.8E-05, R^2^=0.102), and remained significant after excluding AD participants from the analysis. However, the associations were not statistically significant when excluding the *APOE* region and the A/T/N PRS associations did not meet the Bonferroni p-value threshold. LHIPV PRS was also nominally associated with baseline memory (p=0.049, R^2^=0.093).

**Table 3.** PRS associations with baseline memory performance.

|  |  |  |  | No *APOE* | NC/MCI | NC |
| --- | --- | --- | --- | --- | --- | --- |
|  |  |  |  |  | N = 845 | N = 336 |
| PRS Score | Beta | P | Adj R<sup>2</sup> | P | P | P |
| A/T/N | -0.09 | **0.048** | 0.093 | 0.761 | **0.049** | 0.072 |
| Clinical AD | -0.10 | **6.79E-05\*** | 0.102 | 0.107 | **1.58E-05\*** | 0.881 |
| CSF AB<sub>42</sub> | -0.04 | 0.072 | 0.093 | 0.335 | 0.148 | 0.243 |
| CSF Ptau | -0.01 | 0.593 | 0.090 | 0.970 | 0.990 | 0.127 |
| LHIPV | 0.05 | **0.049** | 0.093 | 0.050 | 0.163 | 0.187 |

**Boldface** indicates $P < 0.05$ . \* $P < 1.25E-03$ Bonferroni threshold. N=1,182 unless noted otherwise. No *APOE* denotes PRS excluding *APOE* region results. NC/MCI and NC indicate sensitivity analysis results with the denoted diagnostic groups (as assessed at baseline).

### 3.3 PRS Performance on Longitudinal Cognition

The A/T/N PRS showed the strongest association with longitudinal executive function, where a higher score was associated with a faster decline in executive function, and this association remained statistically significant when removing the *APOE* region or excluding AD participants (**Table 4, Figure 1**; A/T/N p=4.6E-07, R^2^=0.125; A/T/N excluding *APOE* p=2.4E-04, A/T/N excluding AD p=2.3E-07). Clinical AD, CSF Aβ_42_, and CSF ptau PRS models were also significantly associated with longitudinal executive function (Clinical AD p=1.9E-04, R^2^=0.116; CSF Aβ_42_ p=0.005, R^2^=0.118; CSF ptau p=3.6E-04, R^2^=0.117). However, no PRS was associated with change in executive function when restricting the sample to participants with normal cognition at baseline.

**Figure 1.**
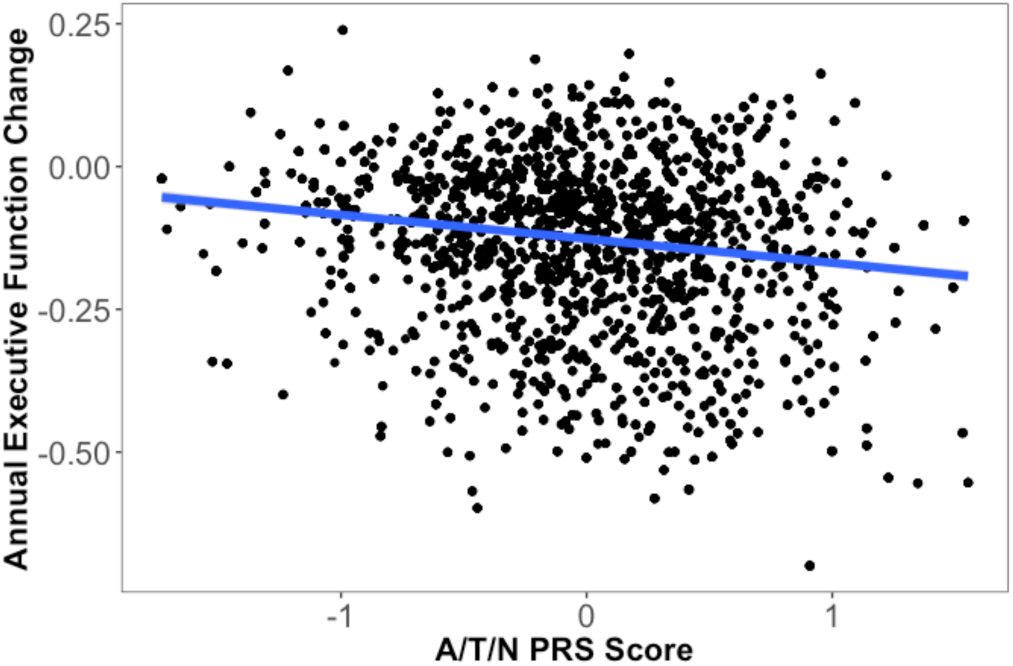
A/T/N PRS score association with annual change in executive function. The A/T/N model accounted for 12.5% of variability in annual executive function change and showed a significant association with longitudinal change in executive function (p=4.62E-07).

**Table 4.** PRS associations with longitudinal executive function performance.

|  |  |  |  | No *APOE* | NC/MCI | NC |
| --- | --- | --- | --- | --- | --- | --- |
|  |  |  |  |  | N = 845 | N = 336 |
| PRS Score | Beta | P | Pseudo R<sup>2</sup> | P | P | P |
| A/T/N | -0.06 | **4.62E-07\*** | 0.125 | **2.40E-04\*** | **2.27E-07\*** | 0.455 |
| Clinical AD | -0.02 | **1.93E-04\*** | 0.116 | 0.083 | **6.90E-05\*** | 0.794 |
| CSF AB<sub>42</sub> | -0.02 | **0.005** | 0.118 | **0.039** | **0.005** | 0.081 |
| CSF Ptau | -0.02 | **3.61E-04\*** | 0.117 | **0.002** | **9.00E-04\*** | 0.689 |
| LHIPV | -0.01 | 0.346 | 0.112 | 0.352 | 0.207 | 0.908 |

**Boldface** indicates $P < 0.05$ . \* $P < 1.25E-03$ Bonferroni threshold. N=1,182 unless noted otherwise. No *APOE* denotes PRS excluding *APOE* region results. NC/MCI and NC indicate sensitivity analysis results with the denoted diagnostic groups (as assessed at baseline).

The A/T/N PRS also showed a significant association with longitudinal memory, where a higher score was associated with a greater rate of memory decline and the association remained statistically significant when removing the *APOE* region or removing AD participants from the analyses (**Table 5, Figure 2**; A/T/N p=1.6E-09, R^2^=0.118; A/T/N excluding *APOE* p=1.2E-05, A/T/N excluding AD p=1.5E-09). Higher clinical AD, CSF Aβ_42_, and CSF ptau PRS were significantly associated with greater decline in memory performance, none of which remained statistically significant when the sample was restricted to individuals with normal cognition at baseline.

**Figure 2.**
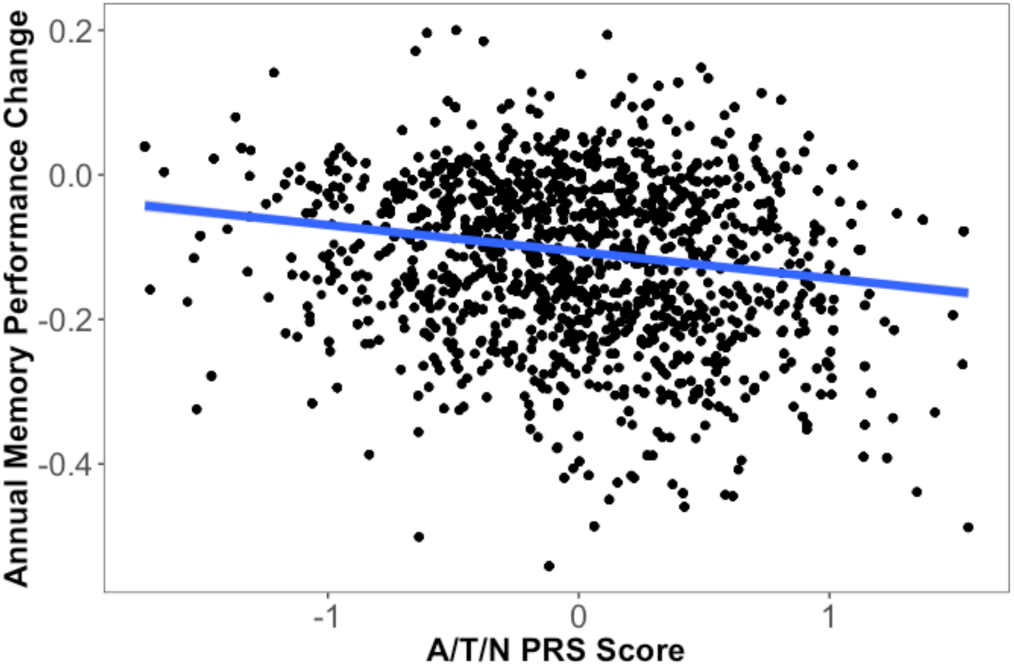
A/T/N PRS score association with annual change in memory performance. The A/T/N model accounted for 11.8% of variability in annual memory performance change, and showed a significant association with longitudinal change in memory performance (p=1.63E-09).

**Table 5.** PRS associations with longitudinal memory performance.

|  |  |  |  | No APOE | NC/MCI | NC |
| --- | --- | --- | --- | --- | --- | --- |
|  |  |  |  |  | N = 845 | N = 336 |
| PRS Score | Beta | P | Pseudo R<sup>2</sup> | P | P | P |
| A/T/N | -0.05 | **1.63E-09\*** | 0.118 | **1.15E-05\*** | **1.53E-09\*** | 0.971 |
| Clinical AD | -0.02 | **1.69E-05\*** | 0.117 | 0.065 | **1.16E-05\*** | 0.474 |
| CSF AB<sub>42</sub> | -0.02 | **1.31E-03** | 0.110 | **0.020** | **1.47E-03** | 0.895 |
| CSF Ptau | -0.02 | **2.55E-04\*** | 0.108 | **1.3E-03** | **1.31E-03** | 0.790 |
| LHIPV | -0.01 | **0.044** | 0.105 | **0.044** | **0.014** | 0.749 |

**Boldface** indicates $P < 0.05$ . \* $P < 1.25E-03$ Bonferroni threshold. N=1,182 unless noted otherwise. No APOE denotes PRS excluding APOE region results. NC/MCI and NC indicate sensitivity analysis results with the denoted diagnostic groups (as assessed at baseline).

In additional sensitivity analyses, we did not observe any sex interactions with PRS on the baseline or longitudinal cognition outcomes detailed above (**Supplemental Table 6**).

## 4. Discussion

We found that the A/T/N PRS with combined genetic risk for CSF ptau_181_, CSF Aβ_42_, LHIPV, and clinical AD was a strong predictor of biomarker levels during the preclinical stages of AD, and a better predictor of longitudinal executive function compared to a PRS for late-onset clinical AD. The A/T/N PRS also performed comparably to the clinical AD PRS on baseline cognitive measures (memory, executive function) and longitudinal memory performance. Together, our findings suggest that utilizing the A/T/N framework by combining genetic risk for AD with genetic risk for AD biomarkers yields a better fitting prediction model, particularly in the earliest stages of disease.

As more GWAS data for AD endophenotypes have become available, it was important that we validate whether PRS for A/T/N are truly predictive of biomarkers in this independent dataset. All PRS were validated for their respective outcomes, and importantly, we found that combining PRS across A/T/N provided a more robust predictor of the individual AD biomarkers. It could be the case that genetic variants with small effects have functional impact across biomarkers, positioning a composite A/T/N score to capture more genetic overlap between biomarkers and amplify relevant signals for associations. As sample sizes continue to increase in genetic studies of amyloid, tau, and hippocampal atrophy in late-life, it is quite likely that the sensitivity of these PRS will continue to improve. Currently, the scores appear to provide a small boost in sensitivity to early deposition of pathology.

Similarly, the A/T/N PRS appeared to provide more sensitivity to risk of executive function decline, particularly when including participants across the clinical spectrum of AD. In contrast, the clinical AD PRS appeared to perform comparably to the A/T/N score in predicting memory performance, suggesting that the current polygenic predictors of clinical AD may be particularly sensitive to memory dysfunction. Given that many of the cohorts included in the AD GWAS studies come from memory clinics across the country, it is not surprising to see such robust associations with memory performance. However, when the *APOE* region is excluded the A/T/N score remains significantly associated with memory decline while the clinical AD score does not, suggesting that the A/T/N score may provide genetic prediction above and beyond the *APOE* region in the context of memory decline. However, the clinical utility of the additional sensitivity provided by the A/T/N PRS in non-memory domains remains to be determined.

It is notable that the associations we observed with biomarkers of neuropathology and longitudinal cognitive decline explained variance above and beyond *APOE*, which has been a limitation of previous PRS analyses [42]. Certainly, *APOE* is a critical genetic component in AD, but AD remains a complex polygenic trait including many genetic effects with small effect sizes. Pooling across these small effects does appear to provide a score that shows more robust associations than reliance on the top genetic signal alone.

Previous work has shown that the clinical AD PRS derived from IGAP summary statistics was predictive of bilateral hippocampal volume in a Brazilian youth cohort [43], aligning with the clinical AD PRS association with left hippocampal volume demonstrated in this study. Together, these results may suggest that genetic drivers of hippocampal volume are influential throughout life. This association also fits with previous literature which reported a PRS for late-onset AD was associated with hippocampal function, as measured by functional magnetic resonance imaging (fMRI) [44]. Few studies to date have tested PRS associations with late life cognitive change, and results have been mixed [45, 46].

A strength of our chosen validation set was the wealth of clinical and genetic data including more than 1,100 participants. However, the GWAS summary statistics and the present analyses were sampled from highly-educated, Non-Hispanic White populations and generalizability of findings to more diverse groups is limited. PRS analyses can capture the common genetic risk for disease, however other disease contributors such as rare variants, gene-gene and gene-environment interactions may not be as accurately modeled. Larger sample sizes for GWAS of endophenotypes in particular would improve power of PRS models. In their current form, PRS models of CSF biomarkers explain a small portion of the variance, suggesting substantial gains could be reached as sample sizes grow.

Overall, this study suggests an integration of genetic components in the A/T/N framework can be used to explain a greater percentage of variability in longitudinal cognition data compared to previously published genetic risk scores for late-onset clinical AD. Importantly, the calculated A/T/N PRS was significantly associated with longitudinal cognition in the absence of the *APOE* region, suggesting a greater degree of genetic risk for cognitive decline can be captured above and beyond *APOE*. Future PRS development should employ endophenotype-specific approaches to predict cognitive trajectories.

