## Supplemental Material for "A/T/N polygenic risk score for cognitive decline in old age"

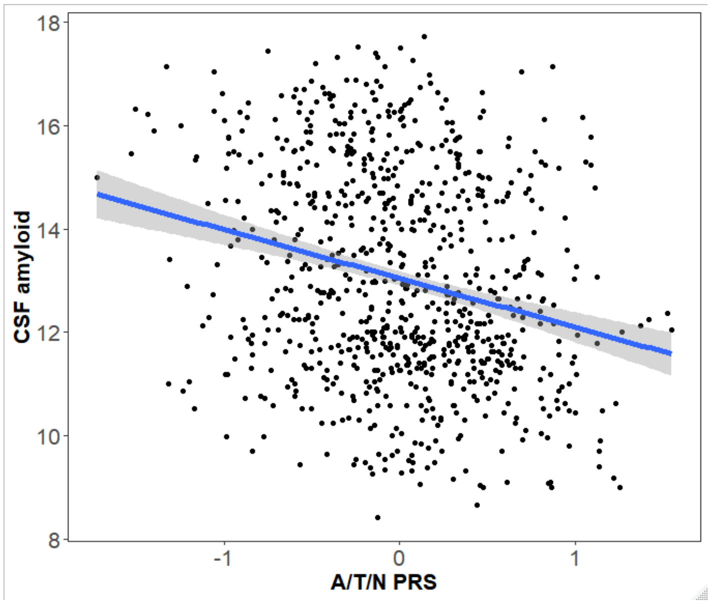


**Supplemental Figure 1.** A/T/N PRS score association with baseline CSF amyloid level. The A/T/N model accounted for 7.6% of variability in the outcome and showed a significant association with baseline CSF amyloid ($p=6.8E-12$).

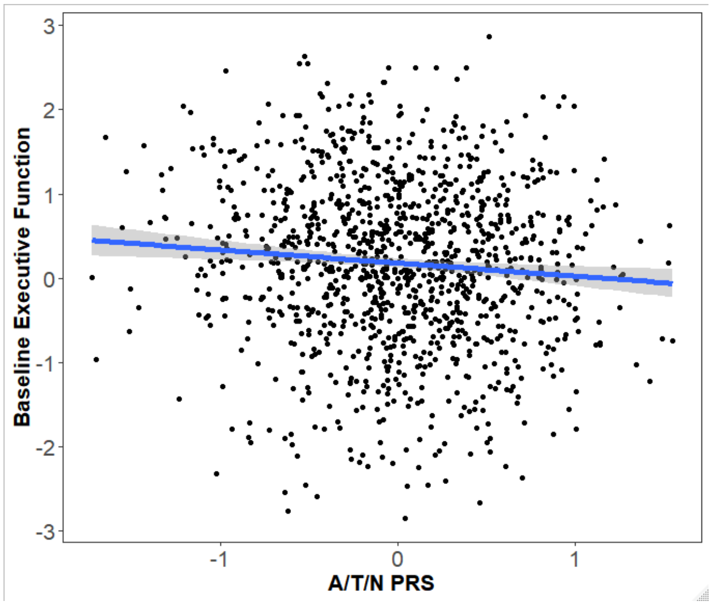


**Supplemental Figure 2.** A/T/N PRS score association with baseline executive function. The A/T/N model accounted for 11.5% of variability in the data and showed a significant association with baseline executive function ($p=0.004$).

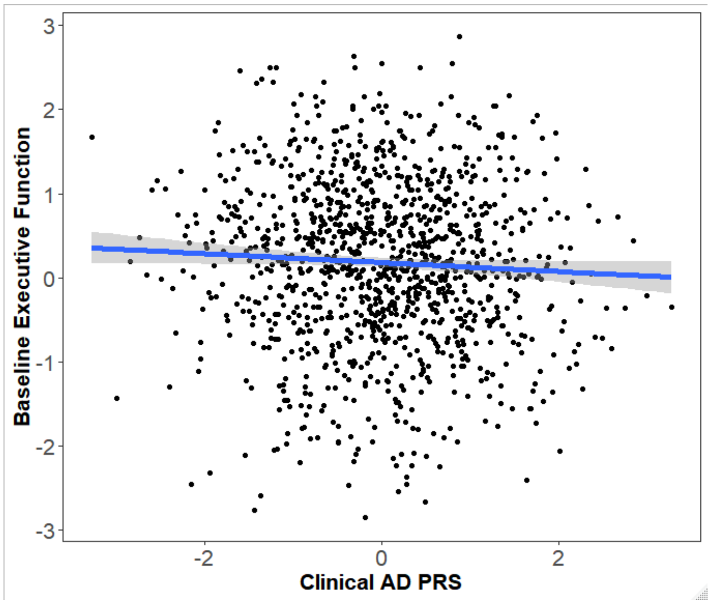


**Supplemental Figure 3.** Clinical AD PRS association with baseline memory performance. The model accounted for 10.2% of variability in the data and showed a significant association with baseline executive function (p=6.8E-05).

**Supplemental Table 1.** Number of SNPs used to calculate PRS

| PRS Score | Number of SNPs used for PRS | Number of SNPs used for PRS excluding *APOE* |
|---|---|---|
| Clinical AD | 7730 | 7645 |
| CSF $AB_{42}$ | 5559 | 5543 |
| CSF Ptau | 6128 | 6118 |
| LHIPV | 8334 | 8329 |

Note: A/T/N score was calculated using the average of scores above.

**Supplemental Table 2**. Model results for baseline CSF amyloid level.

| | | | | No *APOE* | NC/MCI | NC |
|---|---|---|---|---|---|---|
| **PRS Score** | **Beta** | **P** | **Adj $R^2$** | **P** | **P** | **P** |
| **A/T/N** | -0.91 | **6.84E-12*** | 0.076 | **8.20E-06*** | **1.13E-11*** | **1.11E-03*** |
| **Clinical AD** | -0.43 | **3.05E-09*** | 0.062 | **0.015** | **1.32E-08*** | **0.013** |
| **CSF $AB_{42}$** | -0.33 | **4.06E-06*** | 0.046 | **6.46E-04*** | **3.66E-05*** | 0.089 |
| **CSF Ptau** | -0.22 | **0.003** | 0.032 | **0.020** | **0.005** | 0.528 |
| LHIPV | -0.09 | 0.227 | 0.023 | 0.232 | **0.043** | 0.053 |

**Boldface** indicates P < 0.05. *P<1.25E-03 Bonferroni threshold. N=826. Adjusted $R^2$ reported is for the whole model (including covariates). No *APOE* denotes PRS excluding *APOE* region results. NC/MCI and NC indicate sensitivity analysis results with the denoted diagnostic groups (as assessed at baseline).

**Supplemental Table 3**. Model results for baseline CSF Ptau level.

| | | | | No *APOE* | NC/MCI | NC |
|---|---|---|---|---|---|---|
| **PRS Score** | **Beta** | **P** | **Adj $R^2$** | **P** | **P** | **P** |
| **A/T/N** | 0.47 | **8.34E-06*** | 0.051 | **0.003** | **1.02E-06*** | 0.110 |
| **Clinical AD** | 0.17 | **0.003** | 0.038 | 0.412 | **0.003** | 0.801 |
| **CSF $AB_{42}$** | 0.12 | **0.035** | 0.033 | 0.194 | **0.026** | 0.285 |
| **CSF Ptau** | 0.17 | **0.003** | 0.038 | **0.010** | **1.13E-03*** | 0.701 |
| LHIPV | 0.09 | 0.126 | 0.030 | 0.129 | **0.036** | **0.037** |

**Boldface** indicates P < 0.05. *P<1.25E-03 Bonferroni threshold. N=826. Adjusted $R^2$ reported is for the whole model (including covariates). No *APOE* denotes PRS excluding *APOE* region results. NC/MCI and NC indicate sensitivity analysis results with the denoted diagnostic groups (as assessed at baseline).

**Supplemental Table 4**. Model results for baseline left hippocampal volume.

| | | | | No *APOE* | NC/MCI | NC |
|---|---|---|---|---|---|---|
| **PRS Score** | **Beta** | **P** | **Adj $R^2$** | **P** | **P** | **P** |
| A/T/N | -53.45 | 0.074 | 0.216 | 0.645 | **0.017** | 0.983 |
| **Clinical AD** | -59.14 | **2.98E-04*** | 0.223 | 0.074 | **3.09E-05*** | 0.549 |
| **CSF $AB_{42}$** | -35.16 | **0.032** | 0.217 | 0.135 | 0.055 | 0.647 |
| CSF Ptau | -24.65 | 0.136 | 0.215 | 0.293 | 0.118 | 0.945 |
| **LHIPV** | 54.39 | **8.94E-04*** | 0.222 | **8.12E-04*** | **0.017** | 0.306 |

**Boldface** indicates P < 0.05. *P<1.25E-03 Bonferroni threshold. N=1,086. Adjusted $R^2$ reported is for the whole model (including covariates). No *APOE* denotes PRS excluding *APOE* region results. NC/MCI and NC indicate sensitivity analysis results with the denoted diagnostic groups (as assessed at baseline).

**Supplemental Table 5**. Model results for AD diagnosis.

| | | | | No *APOE* |
|---|---|---|---|---|
| **PRS Score** | **Beta** | **Z** | **P** | **P** |
| **A/T/N** | 0.60 | 4.14 | **3.52E-05*** | **0.009** |
| **Clinical AD** | 0.45 | 5.38 | **7.43E-08*** | **1.29E-03** |
| **CSF $AB_{42}$** | 0.21 | 2.81 | **0.005** | **0.048** |
| CSF Ptau | 0.15 | 1.91 | 0.056 | 0.160 |
| LHIPV | -0.07 | -1.00 | 0.319 | 0.318 |

**Boldface** indicates P < 0.05. *P<1.25E-03 Bonferroni threshold. N=657. Adjusted $R^2$ reported is for the whole model (including covariates). No *APOE* denotes PRS excluding *APOE* region results.

**Supplemental Table 6**. No PRS interacted with sex on cognitive outcomes.

| | Baseline EF | Baseline Memory | Longitudinal EF | Longitudinal Memory |
|---|---|---|---|---|
| ***PRS x Sex*** | **P** | **P** | **P** | **P** |
| A/T/N | 0.64 | 0.37 | 0.33 | 0.07 |
| Clinical AD | 0.45 | 0.29 | 0.83 | 0.81 |
| CSF $AB_{42}$ | 0.60 | 0.32 | 0.58 | 0.43 |
| CSF Ptau | 0.36 | 0.74 | 0.28 | 0.07 |
| LHIPV | 0.51 | 0.81 | 0.70 | 0.20 |

EF = executive function. **Boldface** indicates P < 0.05. *P<1.25E-03 Bonferroni threshold. N=1,181.
